# Co-existence of prediction and error signals in electrophysiological responses to natural speech

**DOI:** 10.1101/2020.11.20.391227

**Authors:** Michael P. Broderick, Edmund C. Lalor

**Author notes:** Correspondence: Michael Broderick; Edmund Lalor.

## Abstract

Prior knowledge facilitates perception and allows us to interpret our sensory environment. However, the neural mechanisms underlying this process remain unclear. Theories of predictive coding propose that feedback connections between cortical levels carry predictions about upcoming sensory events whereas feedforward connections carry the error between the prediction and the sensory input. Although predictive coding has gained much ground as a viable mechanism for perception, in the context spoken language comprehension it lacks empirical support using more naturalistic stimuli. In this study, we investigated theories of predictive coding using continuous, everyday speech. EEG recordings from human participants listening to an audiobook were analysed using a 2-stage regression framework. This tested the effect of top-down linguistic information, estimated using computational language models, on the bottom-up encoding of acoustic and phonetic speech features. Our results show enhanced encoding of both semantic predictions and surprising words, based on preceding context. This suggests that signals pertaining to prediction and error units can be observed in the same electrophysiological responses to natural speech. In addition, temporal analysis of these signals reveals support for theories of predictive coding that propose that perception is first biased towards what is expected followed by what is informative.

**Significance Statement:** Over the past two decades, predictive coding has grown in popularity as an explanatory mechanism for perception. However, there has been lack of empirical support for this theory in research studying natural speech comprehension. We address this issue by developing an analysis framework that tests the effects of top-down linguistic information on the auditory encoding of continuous speech. Our results provide evidence for the co-existence of prediction and error signals and support theories of predictive coding using more naturalistic stimuli.

## 1. Introduction

A key question in neuroscience centers on how bottom-up sensory inputs combine with top-down prior knowledge to subserve perception (Miller et al., 1951; Liberman et al., 1967). A popular idea is that this is achieved through hierarchical Bayesian inference, whereby the brain infers the causes of its sensory inputs by using an internal model of the world to generate predictions and then comparing those predictions to incoming sensory input (Knill and Pouget, 2004; Aitchison and Lengyel, 2017). One prominent mechanistic account – known as predictive coding – proposes that higher levels in the cortical hierarchy predict lower-level bottom-up signals via feedback connections, whereas feedforward connection convey only the error between top-down prediction and the bottom up sensory signal (Rao and Ballard, 1999; Friston, 2005; Clark, 2013). A core tenet of this theory assumes the involvement of two distinct populations of neurons at each cortical level: representation (prediction) units, which encode predictions based on prior information, and error units, which encode the error (Rao and Ballard, 1999). Given this assumption, one should expect the co-presence of neural signals reflecting computations from these distinct populations during perception. Indeed, while some evidence exists (Egner et al., 2010), there is an overall lack of neuroimaging studies providing direct empirical support for the simultaneous computation of prediction and prediction error. Thus, there is still ongoing debate as to how prior information informs perception (Egner and Summerfield, 2013; Heilbron and Chait, 2017; Friston, 2018).

Studying speech processing in the brain represents a very powerful way to contribute to this debate. This is because speech perception likely involves predictions across many levels of processing (Kuperberg and Jaeger, 2016), and because the field of computational linguistics has given us methods for quantifying how upcoming words may be predicted from their context (Bengio et al., 2003; Mikolov et al., 2013; Buck et al., 2014; Pennington et al., 2014). In addition, pattern analysis techniques (Haxby, 2001; Kriegeskorte, 2008; Crosse et al., 2016) have helped adjudicate between predictive coding and competing accounts of Bayesian inference by directly testing the stimulus features that are encoded in the neural signal. Studies employing such approaches have produced mixed results, in some case showing evidence of enhanced prediction signals (Kok et al., 2012; Leonard et al., 2016; Broderick et al., 2019) and in other cases showing enhance error signals (Blank and Davis, 2016; Sohoglu and Davis, 2020). The notion of a two-process model where perception is first biased towards prior knowledge and later upweights surprising events (Press et al., 2020) has the potential reconcile these seemingly contrasting findings and also supports the co-existence of representation and error units.

Our current work aims to provide evidence for both representational and prediction error effects in the same responses to natural speech. Specifically, we quantify the predictability of words using measures from language models that have been previously linked to the computation of error and prediction. The first of these measures – known as surprisal – quantifies the Kullach-Leibler divergence between prior and posterior probability distributions and thus reflects the degree of belief updating a comprehender undergoes when processing new incoming words in context (Levy, 2008; Kuperberg and Jaeger, 2016). It has been linked formally with prediction error in theories of predictive coding (Friston, 2005). The second measure – known as semantic similarity – is derived by comparing words to their context based on word embedding models. It has been previously shown to relate to the predictive preactivation of the semantic features of words, something that is thought to underlie the N400 response (Federmeier and Kutas, 1999; Ettinger et al., 2016; Broderick et al., 2020). We assessed how these measures of surprisal and semantic similarity affected the encoding of bottom-up acoustic features using a recently developed analysis framework (Broderick et al., 2019). Importantly, the measures of semantic similarity and surprisal assign higher values to more expected and less expected words, respectively, and are only weakly (and negatively) correlated. However, our results reveal that both measures independently affect the encoding of acoustic-phonetic information. Additionally, we see that representation and error are dissociable based on their timing of their effects, with the representational effect preceding that of the surprisal effect (i.e., the error). This is, again, something that has been hypothesized in the literature (Press et al., 2020).

## 2. Materials and Methods

### 2.1 Participants

Data from 19 native English speakers (6 female; aged 19-38 years) who participated in a previous study was reanalysed for this study (Di Liberto et al., 2015; Broderick et al., 2018). The study was undertaken in accordance with the Declaration of Helsinki and was approved by the Ethics Committee of the School of Psychology at Trinity College Dublin. Each subject provided written informed consent. Subjects reported no history of hearing impairment or neurological disorder.

### 2.2 Stimuli and experimental procedure

Subjects undertook 20 trials, each just under 180 seconds in length, where they were presented with an audio-book version of a popular mid-20th century American work of fiction (Hemingway, 1952), read by a single male American speaker. The average speech rate was 210 words/minute. The mean length of each content word was 340 ms with standard deviation of 127 ms. Trials were presented chronologically to the story. All stimuli were presented diotically at a sampling rate of 44.1 kHz using Sennheiser HD650 headphones and Presentation software from Neurobehavioural Systems. Testing was carried out in a dark, sound attenuated room and subjects were instructed to maintain visual fixation on a crosshair centred on the screen for the duration of each trial, and to minimise eye blinking and all other motor activities.

### 2.3 EEG acquisition and preprocessing

128-channel EEG data were acquired at a rate of 512 Hz using an ActiveTwo system (BioSemi). Triggers indicating the start of each trial were sent by the stimulus presentation computer and included in the EEG recordings to ensure synchronization. Offline, the data were bandpass filtered between 1 and 8Hz using a Chebyshev Type II filter (order 54, cutoff 0.5Hz for high pass filtering and 8.5Hz for low pass filtering). Passband attenuation was set to 1dB and stopband attenuation was set to 60dB (high pass) and 80dB (low pass). After filtering, data were downsampled to 64 Hz (backward modelling) or 128 Hz (forward modelling; see below). To identify channels with excessive noise, the standard deviation of the time series of each channel was compared with that of the surrounding channels. For each trial, a channel was identified as noisy if its standard deviation was more than 2.5 times the mean standard deviation of all other channels or less than the mean standard deviation of all other channels divided by 2.5. Channels contaminated by noise were recalculated by spline interpolating the surrounding clean channels in EEGLAB (Delorme and Makeig, 2004). The data were then re-referenced to the global average of all channels.

### 2.4 Stimulus characterisation

Several acoustic and linguistic features were extracted from the speech signal and used as input at various stages in a 2-stage regression analysis. These features can be categorised into 3 main groups based on what stage they are used in the analysis.

1. The first group of features were regressed to the recorded EEG signal using the temporal response function (see below), which learns a linear mapping between stimulus and neural response or vice versa. Each feature was used to assess the encoding of speech at different hierarchical levels.
  1a. *Envelope*. The broadband amplitude envelope of the speech signal was calculated using the absolute value of the Hilbert transform.
  1b. *Spectrogram*. The speech signal was filtered into 16 different frequency bands between 250Hz and 8kHz according to Greenwood’s equations (Greenwood, 1961). After filtering, the amplitude envelope was calculated for each band using the absolute value of the Hilbert Transform.
  1c. *Phonetic Features*. To create the phonetic feature stimulus Prosodylab-Aligner software (Gorman et al., 2011) was used. This automatically partitions each word in the story into phonemes from the American English International Phonetic Alphabet (IPA) and performs forced-alignment, returning the starting and ending time-points for each phoneme. Each phoneme was then mapped to a corresponding set of 19 phonetic features, which was based on the University of Iowa’s phonetics project http://prosodylab.cs.mcgill.ca/tools/aligner/.
2. The second group of features estimates higher-level linguistic properties of words and was used in the second stage of the regression analysis.
  2a. *Semantic Similarity* was calculated for each content word in the narrative. It is estimated from word embeddings derived using GloVe (Pennington et al., 2014) which was trained on a large corpus (Common Crawl https://commoncrawl.org/). Each word is represented as 300-dimensional vector where each dimension can be thought to reflect some latent linguistic context. A word’s similarity index is estimated as the Pearson’s correlation between the word’s vector and the averaged vector of all the preceding words in the sentence (Broderick et al., 2018). Thus, small similarity values signify out-of-context words that, by extension, are unexpected.
  2b. *Surprisal* was calculated using a Markov model trained on the same corpus as GloVe (Common Crawl). These models, commonly referred to as n-grams, estimate the conditional probability of the next word in a sequence given the previous n-1 words. We applied a 5-gram model that was produced using interpolated modified Kneser-Ney smoothing (Chen and Goodman, 1996; Buck et al., 2014). Unlike semantic similarity, surprisal assigns higher values to more unexpected words.
3. The third group of stimulus characterisations was also used in the 2nd stage regression analysis to act as nuisance regressors. Their purpose was to soak up any variance in word level auditory encoding relating to acoustic changes in the speaker’s voice.
  3a. *Envelope Variability* was calculated by taking the standard deviation of the speech envelope over each spoken word. Here, we wished to control for rapid changes in the envelope amplitude as it has been shown that cortical responses monotonically increase with steeper acoustic edges (Oganian and Chang, 2019).
  3b. *Relative Pitch* was recently shown to be encoded in EEG (Teoh et al., 2019). It quantifies pitch normalised according to the vocal range of the speaker. Praat software (Boersma and Weenink, 2000) was used to extract a continuous measure of pitch (absolute pitch). The measure was then normalized to zero mean and unit standard deviation (z-units) to obtain relative pitch.
  3c. *Resolvability* measures whether the harmonics of a sound can be processed within distinct filters of the cochlea (resolved) or if they interact within the same filter (unresolved). It has previously been shown using fMRI that pitch responses in auditory cortex are predominantly driven by resolved frequency components (Norman-Haignere et al., 2013). Custom written scripts from an acoustic statistics toolbox from the same study were used to extract a continuous measure of harmonic resolvability.

### 2.5 Evaluating cortical encoding at the word level

Our analysis framework consists of two stages of regression (Figure 1). The first stage models the relationship between audio stimulus and neural response, thus quantifying how well certain features of speech were encoded in the neural signal. This was done using linear regression in which two separate models were trained: a backwards model, which reconstructs features of the stimulus from the neural response, and a forward model, which predicts neural responses from the stimulus.

**Figure 1.**
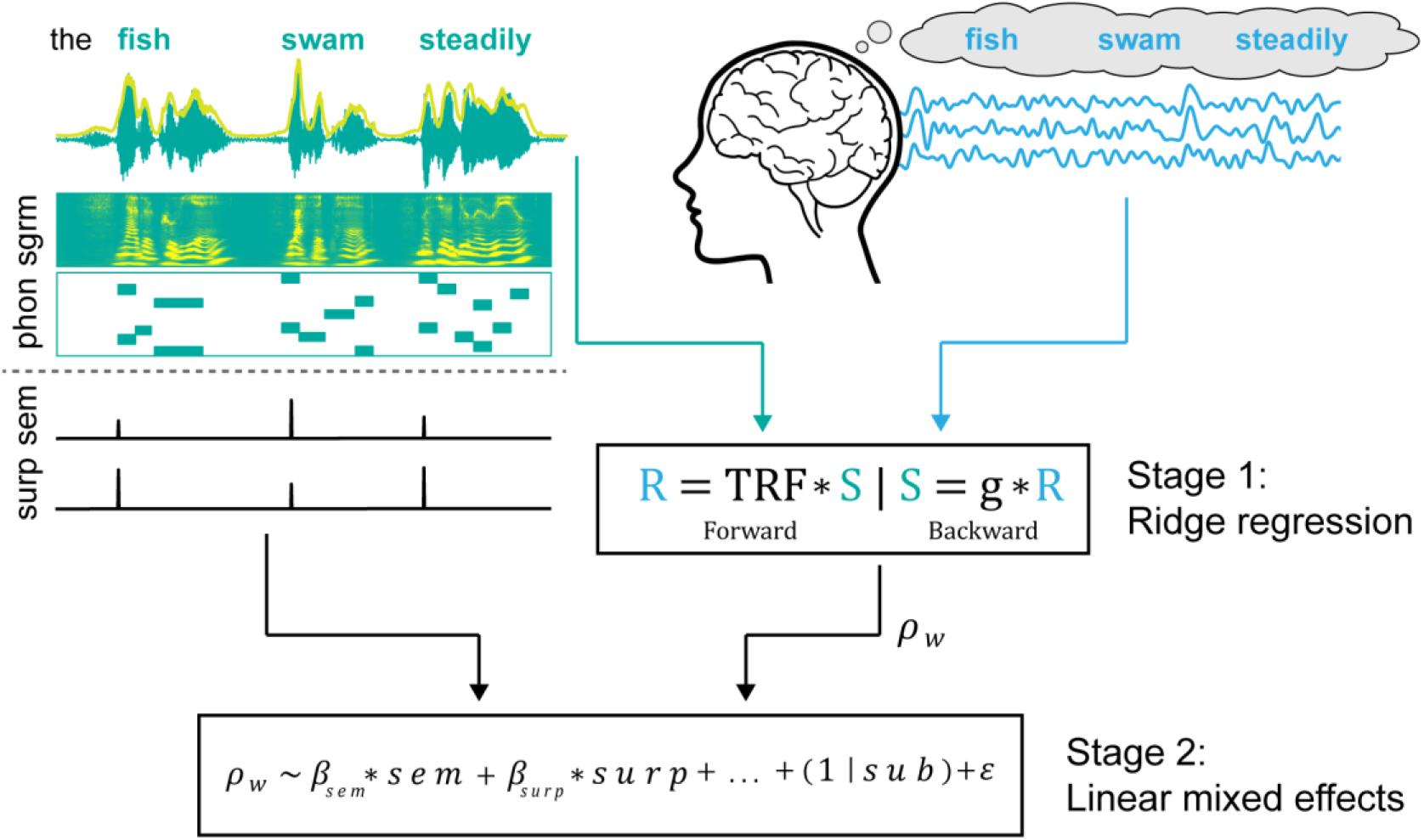
Analysis framework. A 2-stage regression analysis framework was used to evaluate the impact of top-down linguistic features on the neural encoding of spoken words. Stage 1 measures the relationship between stimulus and response using ridge regression. A backwards TRF was used to reconstruct the speech envelope and a forwards TRF was used to predict EEG based on the acoustic and phonetic features of speech. This resulted in a set of reconstruction/prediction accuracies which were used as dependent variables in the 2^nd^ stage model. Stage 2 measures the relationship between the encoding of individual words’ speech features and lexico-semantic properties. Semantic similarity and surprisal are estimated for each word using computational language models. A linear mixed effects model is used to assess the relationship between these features and reconstruction/prediction accuracies.

For the backwards modelling approach, a backwards TRF or decoder was trained to reconstruct an estimate of the speech envelope from the neural data (Mesgarani et al., 2009; Ding and Simon, 2012). A decoder, g_n_(τ), represents the linear mapping from neural response, r_n_(t), to stimulus, s(t), expressed by the following equation

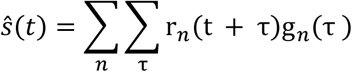

where 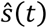 is the reconstructed stimulus envelope; n represents the recorded neural channels and τ is a specified number of time lags ranging from −100 to 300ms. Decoder weights were estimated using regularized linear regression, wherein a regularization (ridge) parameter, λ, was tuned to control overfitting. In a leave-one-out cross-validation procedure, a decoder, trained on all but one trial, was used to reconstruct an estimate of the speech envelope of the left-out trial.

The forward modelling of stimulus to response was done using the Temporal Response Function (TRF) (Lalor and Foxe, 2010). For N recorded channels the instantaneous neural response r(t, n), sampled at times t =1….T and channel n, consists of a convolution of the stimulus property, s(t), with a channel specific TRF, w(τ, n). The response can be modelled as

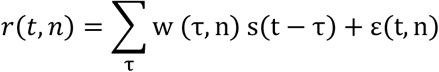

TRF weights are also estimated using regularized linear regression and a leave-one-out cross-validation procedure. TRFs, trained on all but one trial, are used to predict EEG response at each channel. This predicted response is compared with the recorded EEG response to assess the model’s ability to capture the encoding of the stimulus feature. Three separate forward TRFs were used based on representing the speech input as its spectrogram (spectrogram-only), a discrete set of time-aligned phonetic features (phonetic features-only) and as a combination of its spectrogram and phonetic features. These models were used to generate three separate sets of EEG predictions that reflect cortical responses to the spectral and phonological properties of speech.

The brain’s encoding of individual words, *ρ*_*w*_, was evaluated in two ways: 1) using a trained decoder to reconstruct the speech envelope of each word in a held-out trial and comparing that with the actual word’s speech envelope, and 2) using a trained TRF to predict the channel-specific EEG response to each word in a held-out trial and comparing that with the actual EEG response at each channel. The comparison was done by measuring the Spearman’s correlation between the two signals for the first 100ms after word onset.

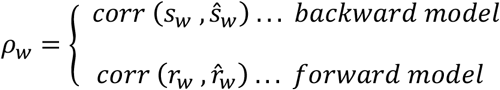

Thus, encoding of individual words was quantified by their reconstruction/prediction accuracy. This was repeated for every trial in the dataset resulting in reconstruction/prediction accuracies for all content words in the experiment. There were, on average, 2 phonemes in every 100ms window. See Crosse et al., 2016 and Broderick et al., 2019 for more details on model training and evaluation.

### 2.6 Measuring the effect of top-down linguistic information and encoding of words

The prediction or reconstruction accuracy at the level of individual words was passed to stage 2 of the analysis framework, where a linear mixed effects (LME) model measured the relationship between the encoding of individual words and their semantic dissimilarity and surprisal relative to preceding context. Here, word-level (reconstruction/prediction) accuracy was used as the dependent variable in the model whereas semantic similarity and surprisal were used as predictor variables. LME models variability due to items and subjects simultaneously. The model is described below.

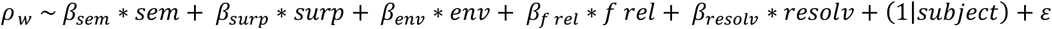

In addition to semantic similarity (*sem*) and surprisal (*surp*), low level acoustic correlates of pronunciation, measured in the first 100ms after word onset, were included in the model as nuisance regressors to ensure that any effect of semantic information on cortical tracking was not in fact an indirect one, by proxy of changes in the speaker’s voice rather than top-down effects in the listener’s brain (Lieberman, 1963). These were envelope variability (*env*), average relative pitch (*f rel*) and average resolvability (*resolv*). All variables were normalized to zero mean and unit standard deviation (z-units) before being input to the model. Pairwise correlations for the predictor variables in the 2nd stage regression were calculated using Pearson’s correlation and are provided in Table 1.

**Table 1.**
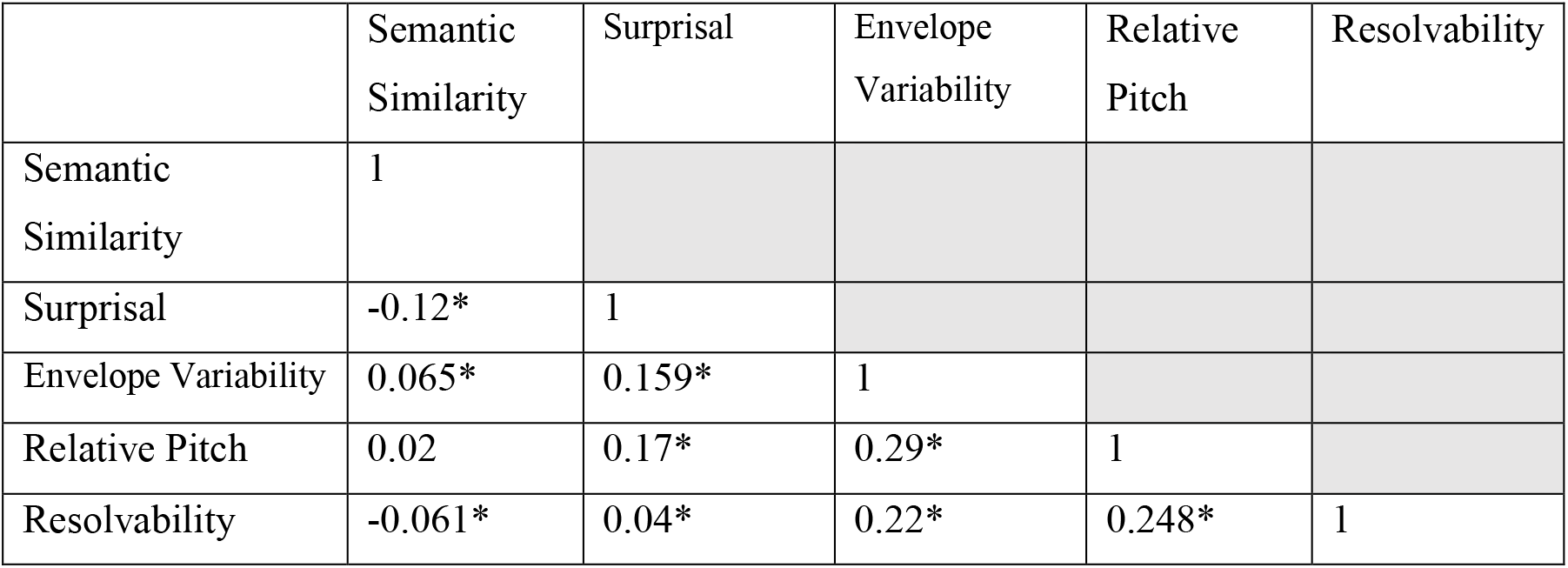
Pairwise correlations between predictors variable in 2^nd^ stage regression (analysis 2). Asterisks indicate significant correlation after correcting for multiple comparisons.

For the forward modelling analysis, EEG prediction accuracies, derived from the combined spectrogram and phonetic features TRF, were used as the dependent variable in the 2nd stage linear mixed effects model, with identical predictor variables as the 2nd stage backwards model. To investigate the top-down effects on isolated measures of acoustic and phonological processing (as opposed to their combination), two 2nd stage models were constructed. These models included prediction accuracies from the spectrogram-only TRF or the phonetic features-only TRF as additional nuisance regressors for each other. The rationale for including these nuisance regressors was that phonological feature and spectrogram representations of speech share redundant information. So, for example, including EEG prediction accuracies from the spectrogram TRF allowed us to partial out that redundant contribution and identify the effects of semantic information on phonetic feature processing in isolation. The analogous analysis was carried out to identify effects of semantic context on spectrographic processing in isolation also.

### 2.7 Statistical analysis

#### Permutation Testing

To test the significance of our model, we ran repeated permutation tests in which reconstruction or prediction accuracy values were fixed and semantic similarity values were randomly shuffled between words, taking the parameters of the model at each permutation. The testing consisted of 500 permutations. Parameters of the true model (β) were deemed significant if they exceeded the 95th percentile of the distributions of model parameters based on shuffled semantic similarity values.

#### Correcting for Multiple Comparisons

Multiple comparison correction was required for analyses that involved statistical testing at multiple scalp electrodes, multiple time windows or both. A cluster-mass non-parametric analysis was conducted (Maris and Oostenveld, 2007) to overcome the multiple comparison problem. This approach includes biophysically motivated constraints that can increase the sensitivity of the statistical test in comparison to a standard Bonferroni correction. Neighbouring electrodes or time windows that show a significant correlation between semantic similarity and reconstruction/prediction accuracy (measured as the T value of the regression coefficient being higher than the critical threshold value; α = 0.05) were clustered together. Cluster-level statistics were calculated by summing the T-values in each cluster separately. Permutation testing was then performed on all the data and Monte Carlo p-values were calculated for all significant clusters under the permutation distribution of the maximum cluster-level statistic (Maris and Oostenveld, 2007).

## 3. Results

### 3.1 Dissociable effects of surprisal and semantic similarity on encoding of low-level speech features

We first investigated the impact of surprisal and semantic similarity on the neural encoding of the speech envelope. This was done using a 2-stage regression approach where stage 1 included decoding the speech envelope from the neural data using a backward model. Decoding accuracy of individual words as well as the higher-level linguistic measures were passed to a 2^nd^ stage linear mixed effects model. The coefficients or beta weights of this model show a significant positive relationship between a word’s semantic similarity to its context and how well its low-level auditory features were encoded (β = 0.017, t = 7.99, p =1.4 × 10^−15^). In addition, there was also a significant positive relationship between how surprising a word was relative to its context (i.e., its surprisal) and how well it was encoded in the brain (β =0.009, t = 4.29, p = 1.7 × 10^−5^). This finding was striking given the fact that the linguistic measures reflect opposite patterns of words expectancy, with semantic similarity assigning higher values to more expected words and surprisal assigning higher values to less expected words. Indeed, this difference was confirmed statistically, with the measures being negatively correlated with each other (Pearson’s r = −0.12, p = 2.18 × 10^−18^). Importantly, both surprisal and semantic similarity were significant despite the inclusion of several nuisance regressors aimed at controlling for acoustic properties of the speech signal. Indeed, these nuisance regressors also showed significant beta weights with respect to envelope reconstruction accuracies (Envelope variability: β = 0.063, p =2.4 × 10^−168^, Relative Pitch: β = −0.017, p = 2.5 × 10^−8^, Resolvability: β = −0.023, p =6.2 × 10^−24^) which was unsurprising given that cortical responses are sensitive to such acoustic measures (Oganian and Chang, 2019). Despite, these measure accounting for a substantial amount of the variance explained in envelope reconstruction accuracy, both surprisal and semantic similarity remained significant.

In addition to the LME model, we ran 2^nd^ stage linear models for each subject individually to obtain single subject beta coefficients. These models were based on the word reconstruction accuracies obtained for each subject separately. Figure 2A shows the distribution model coefficients of surprisal and semantic similarity. These weights were both significantly greater than zero (t_18_ = 6.75, p = 2.5 × 10^−6^ for similarity and t_18_ = 4.07, p = 7.21 × 10^−4^ for surprisal). We conducted permutation testing, shuffling either semantic similarity or surprisal predictor variables while holding the other predictor variable constant. The beta coefficients of the shuffled parameter were extracted from each subject’s individual model with each permutation. Overlaid histograms in figure 2A show the distribution of median values across subjects for each shuffled permutation. The median values for the unshuffled models (boxplot line) exceed the highest median values of the permuted models.

**Figure 2.**
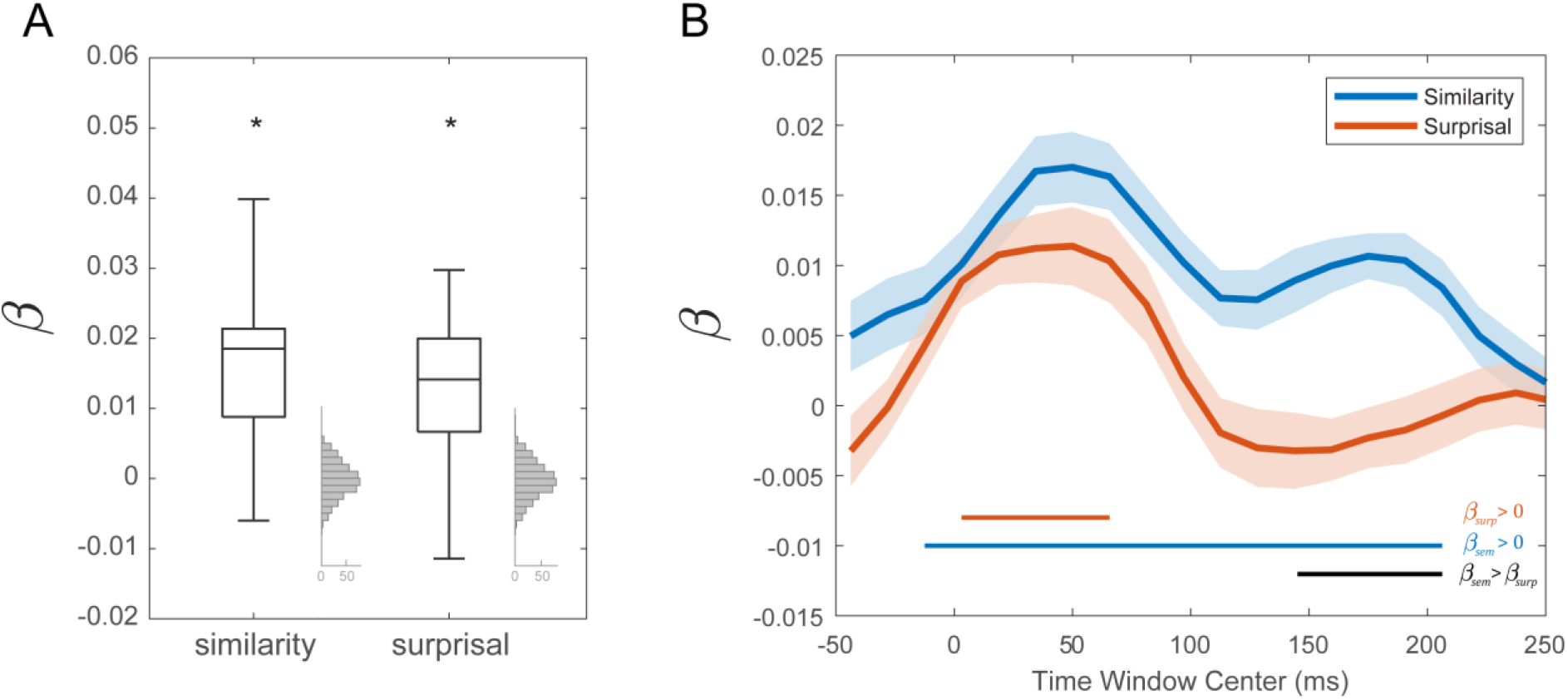
Surprisal and semantic similarity β coefficients for 2^nd^ stage model. For the 2^nd^ stage model, semantic similarity and surprisal were regressed against word reconstruction accuracies outputted from the 1^st^ stage backwards TRF model. Separate models were estimated for each individual subjects **(A)** Box-and-whisker plots of sematic similarity and surprisal β coefficients of 2^nd^ stage models for each subject. Overlaid histograms show the distribution of median values across subjects for each shuffled permutation. The median values for the unshuffled models (box mid-line) exceed the highest values of the permuted models. **(B)** Semantic similarity and surprisal β coefficients as a function of the time window used for estimating word reconstruction accuracy. β coefficients were averaged across subjects, shaded area indicates s.e.m. Both coefficients were significantly greater than 0 (blue and red horizontal line, FDR corrected) in earlier windows after word onset. However, β coefficients for semantic similarity showed sustained significant values at later time windows. Semantic similarity β coefficients were significantly greater than surprisal β coefficients in later time windows (black horizontal line, FDR corrected).

The reconstruction accuracies for each word were based on the correlations between the envelope and the reconstructed envelope in the first 100ms after word onset. To study how top-down information affects the encoding of a word as it unfolds in time, we shifted this 100ms window over the word, recalculating reconstruction accuracies at each 15ms increment. Each set of reconstruction accuracies was regressed with the fixed lexico-semantic measures to calculate a model weights for each time window. Figure 2B shows the resulting weight values, averages across subjects. Beta weights for both similarity and surprisal show peaks in an early time window of ~0-100ms and were both significant in this window (p < 0.05, running t-test, FDR corrected). In addition, beta weights for semantic similarity showed a sustained significant positivity that were significantly greater than those for surprisal at a later time window of ~100-250ms (p < 0.05, running paired t-test, FDR corrected).

### 3.2 Phonetic encoding of speech reflects prediction and prediction error at different latencies

The encoding of acoustic and phonetic speech features was further assessed using the forward TRF. This method predicts channel specific neural responses from the speech input and thus provides information on scalp locations where interactions between high-level semantic information and low-level auditory processing are most prominent. It also enables analysis based on representing speech in multi-dimensional feature spaces, which allows for the disentangling of contributions of different speech features that are likely conflated in the single univariate envelope measure. Previous work investigating the impact of semantic similarity on the processing of spectrotemporal and phonetic feature representations of speech has shown that distinct patterns emerge when the encoding of these features are viewed in isolation (Di Liberto et al., 2015, 2018; Prinsloo and Lalor, 2020; Teoh and Lalor, 2020). TRFs were trained on 1) a combined spectrogram and phonetic feature speech representation, 2) a spectrogram representation and 3) a phonetic feature representation. Prediction accuracies from the 2) spectrogram TRF or 3) the phonetic feature TRF were include as nuisance regressor in the 2^nd^ stage model of the 1) combined TRF in order to isolate responses of the reciprocal feature. Similar to the backward modelling approach, β was estimated as function of time by incrementally shifting the 100ms window that calculated word-level prediction accuracies.

Figure 3A-B shows the semantic similarity and surprisal β weights as a function of time window centre for 2^nd^ stage isolated spectrogram and phonetic feature models. Weights were averaged across subjects and fronto-central electrodes. Figure 3C-D show β weights for semantic similarity and surprisal at each EEG channel for two selected, non-overlapping time windows of 0-100ms and 100-200ms (window centres of 50ms and 150ms respectively). Black dots indicate the channels and time windows where beta coefficients were significantly greater than zero, after correcting for multiple comparisons. There was a lack of effect of higher-level linguistic features of words on the encoding of their spectrographic properties as index by the β weights for the isolated spectrogram model. However, we observed a cluster of right temporal electrodes where β was significant. In contrast, we observed a strong relationship between the lexico-semantic measures and the phonological processing of speech. Beta weights for surprisal and semantic similarity were significant over fronto-central electrodes. Furthermore, the influence of these measures on bottom up processing was dissociated in time, with semantic similarity exerting a stronger influence at the earlier time window and surprisal exerting a stronger influence in the later window. A 2-way ANOVA with factors of time window (0-100ms and 100-200ms) and measure (similarity and surprisal) further supported this finding, revealing a significant interaction (F = 5.93, p < 0.02; figure 3F) and no main effects (F = 0.01, p > 0.05 and F = 0.1, p > 0.05, for measure and time-window effects, respectively). Similarity and surprisal weights were compared in each time window separately, post hoc. Similarity beta weights were significantly higher than surprisal beta weights in the earlier time window (t_18_ = 2.37, p = 0.008, paired t-test) and surprisal beta weights were significantly higher than similarity weight in the later time window (t_18_ = 2.18, p = 0.005, paired t-test). We observed no significant main effects or interaction for the spectrogram model (F = 0.09, p > 0.05, F = 0.83, p > 0.05 and F = 0, p > 0.05 for measure, time-window and interaction effects, respectively).

**Figure 3.**
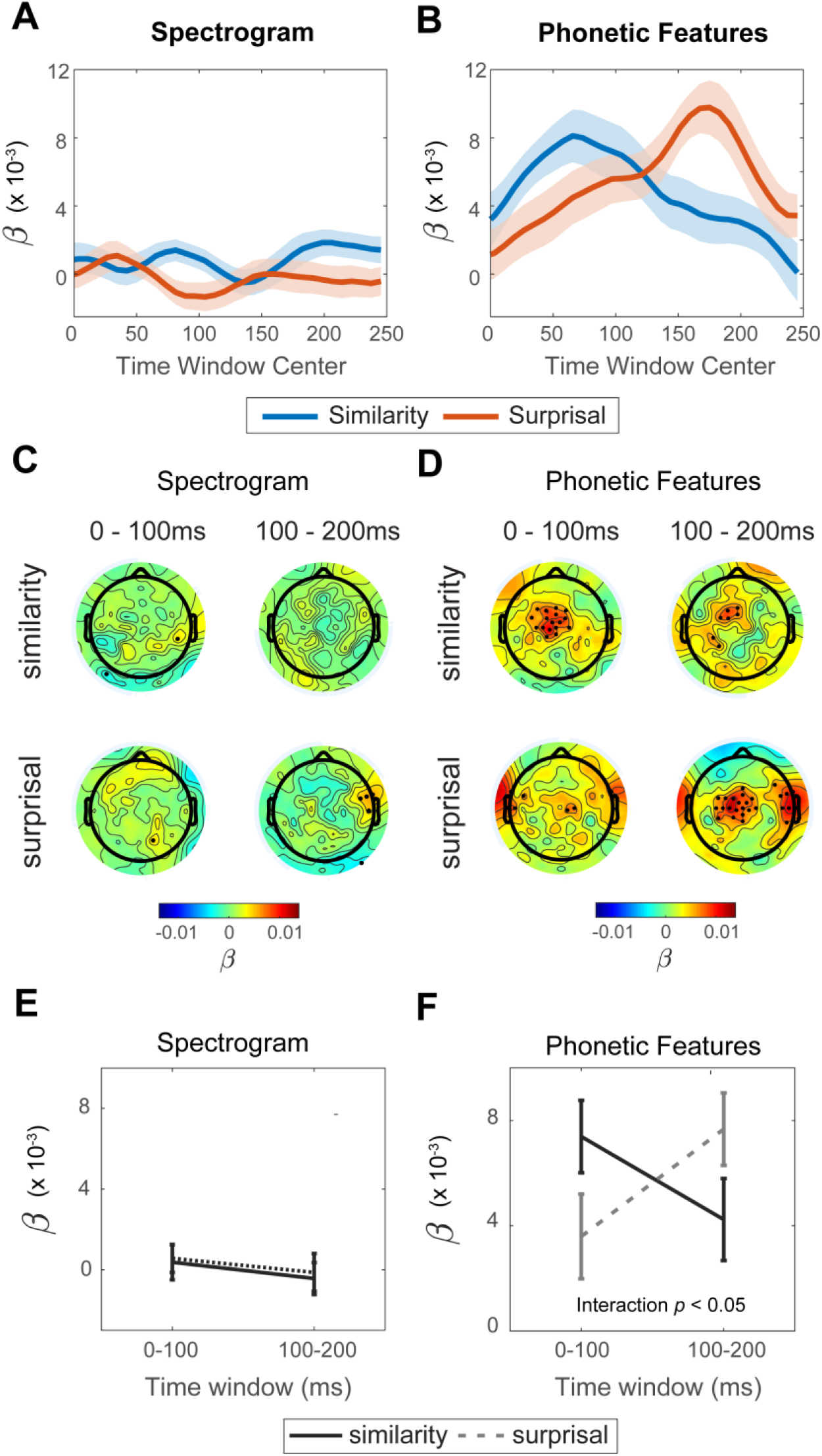
β coefficients for 2^nd^ stage model based on the forward TRFs. **(A-B)** β coefficients for surprisal and semantic similarity averaged over fronto-central EEG channels and across subjects plotted as a function of time window for **(A)** isolated spectrogram and **(B)** isolated phonetic features. Shaded areas indicate s.e.m. **(C-D)** Topographical plots show LME model coefficients for semantic similarity and surprisal as a function EEG channel and time window. Black dots indicate the channels and time points where the β was significantly greater than 0, after multiple-comparison correction. β coefficient for phonetic features show strong activation. In contrast significant β coefficients for the spectrogram model are largely absent. **(E-F)** Average weight coefficients (± s.e.m.) of individual subject 2^nd^ stage models over frontal central electrodes as a function of time window. A significant interaction between lexico-semantic measure and time-window was found for the phonetic feature model.

## 4. Discussion

Previous work analysing EEG responses to natural speech (Broderick et al., 2019) has supported the view that prior knowledge about upcoming words influences their perception at lower representational levels (McClelland and Elman, 1986; McClelland et al., 2006). However, there are different ways in which this top-down effect can be computationally realised (de Lange et al., 2018); and different models based on Bayesian inference have sought to explain precisely how top-down information combines with bottom up sensory processing to subserve perception (Aitchison and Lengyel, 2017). Predictive coding theories argue that only the discrepancies between predictions and true sensory events are propagated up the cortical hierarchy through feedforward connections (Rao and Ballard, 1999; Friston, 2005; Clark, 2013). However, in context of everyday spoken language evidence for neural signatures relating to both prediction and error has been lacking. In this study, we show that both highly surprising words and words whose semantic contents is highly predictable are more strongly encoded by the brain in the early stages of their utterance.

Alternative theories to predictive coding, known as sharpening, argue that the encoding of predicted stimulus features is enhanced by both feedforward and feedback connections (Lee and Mumford, 2003; Kok et al., 2012). Adjudicating between sharpening and predictive coding theories has been difficult using analyses that compare neural response amplitudes, as both theories predict a reduction in neural activity for more expected stimuli (Kersten et al., 2004). Techniques like linear regression, representational similarity analysis (RSA) and multivoxel pattern analysis (MVPA) that more directly measure the patterns encoded in neural activity have provided a breakthrough in this regard (Haxby, 2001; Kriegeskorte, 2008). That said, studies using such techniques have yielded apparently contradictory findings (Kok et al., 2012; Blank and Davis, 2016; Blank et al., 2018; Wang et al., 2018; Sohoglu and Davis, 2020). Here we have shown that combining linear regression methods with linguistic features derived from language models that reflect the predictability (semantic similarity) and surprisal of words supports theories of predictive coding, where signatures of prediction and error can be detected in the same neural signal.

Semantic similarity is derived using word embedding models that capture the latent semantic features of words. It compares a word’s semantic features with those that been activated by the context in which it appears. We propose that this measure reflects a neural process in which semantic features of upcoming words are predictively preactivated ahead of their perception (Federmeier and Kutas, 1999; Kuperberg and Jaeger, 2016). The more strongly a word’s context preactivates features that match its own, the more predictable that word will be at the level of semantics. Importantly, predictions at these higher levels propagate down the hierarchy to activate internal phonetic and acoustic representations before the arrival of bottom up information (Van Petten et al., 1999; DeLong et al., 2005). We show enhanced neural encoding of speech features for words with higher semantic similarity to their previous context. Such effects align with findings by Leonard and colleagues. They show that, when a listener is biased by semantic context, they perceive ambiguous sounds as certain phonemes that conform to that bias (Leonard et al., 2016). Crucially, this perception is underpinned by an enhanced neural representation of the perceived phoneme, despite it being omitted from the bottom up signal. This indicates that top down semantic information penetrates perceptual processing at the level of phonemes. Previous studies have observed a similar enhanced neural representation of predictable events (Kok et al., 2012) and have taken this as evidence for the sharpening of predicted signals. On its own, our observed positive relationship between semantic similarity and neural encoding would support similar conclusions. However, using a measure of Bayesian surprise that relates more directly to prediction error expands on this finding (Friston, 2005; Levy, 2008).

Bayesian surprise reflects the degree of belief updating that a comprehender undergoes as they shift from a prior to posterior distribution of beliefs. As discussed above, it is one way of computationally formalising prediction error (Friston, 2005; Levy, 2008). In contrast to semantic similarity, we found that words that were more surprising (less expected) were more strongly encoded in the neural signal. Previous experiments have observed similar effects, with stronger encoding of more unexpected sensory signals (Gagnepain et al., 2012; Blank and Davis, 2016; Sohoglu and Davis, 2020). These findings, as well as our own, support theories of predictive coding. Crucially though, our analysis revealed the parallel computations of prediction and prediction error.

We employed two separate modelling approaches within our analysis framework. Using the backward model, word level envelope reconstruction accuracies measured how well words were encoded by the brain during the time of their utterance. The forward model similarly indexed the strength of word level encoding but allowed for this encoding to be examined at more fine-grained levels of representation. Importantly, both approaches revealed a significant effect of surprisal and semantic similarity on the early auditory encoding of words. However, differences emerged in the strength and timing of these effects, depending on the type of representation and the modelling approach used in the analysis. Using the forward model, we observed that lexico-semantic features have a stronger influence on the encoding of speech at the level of phonetic features than at the level of the spectrogram. This could be due to phonetic features being a more invariant representation of the speech stimulus (Chang et al., 2010). Thus, the representation corresponds more directly to *which* particular words or phonemes are being spoken at any given time, rather than *how* they vary acoustically from utterance to utterance, as captured by the spectrogram representation. During language comprehension, although the brain may be engaged in predicting both components of speech, it is reasonable to think that what is being spoken will be more strongly predicted and represented in the EEG signal (Kutas and Hillyard, 1980). A difference in the timing of β for semantic similarity and surprisal also emerged in the forward model that was absent in the backward model. Using the forward modelling approach to test the top-down effects phonetic feature encoding we observed earlier effects of semantic similarity before effect of surprisal on cortical encoding. In contrast, using the backwards modelling approach, β coefficients for semantic similarity and surprisal had a similar early time course. This could be due to the speech representations being used at the 1^st^ stage of the model. The speech envelope represents the amplitude fluctuations in the speech signal and envelope tracking likely reflects conflated activity from the encoding of speech at multiple representational levels (Brodbeck and Simon, 2020). In contrast with the forward modelling approach, acoustic factors were controlled for in the 2^nd^ stage regression to isolate the encoding of speech at the phonetic level. Efforts continue aimed at disentangling the different contributions to envelope encoding (Oganian and Chang, 2019; Prinsloo and Lalor, 2020).

The dissociation in time between semantic similarity and surprisal β coefficients gives compelling evidence in favour of predictive coding and conforms with intuitions underlying this theory. The co-existence of distinct populations at each cortical level that encode predictions and prediction errors is a core tenet of predictive coding theories which has been questioned due the apparent lack of empirical support (Egner and Summerfield, 2013). Our results indicate the coexistence of these units and show them in operation during the processing of naturalistic continuous speech. The use of recordings with high temporal resolution like EEG allows us to observe the timings of these operations in finer detail and produces results that further align with this view. We observed an interaction with the lexico-semantic measures of prediction and the relative timings of speech feature enhancement. Neural processing of features is first enhanced for more expected words, followed by the enhancement of words that are more surprising. This result conforms to the intuition of predictive coding. The brain must first capture predictions about an upcoming signal before the error can be computed (Press et al., 2020).

The idea that semantic similarity and surprisal map directly to representation and error units, respectively, is by no means conclusive. Indeed, predictions about upcoming words as indexed with surprisal could be captured in representation units and vice versa. However, our results suggest that these measures, on average, reflect response properties in these distinct units. The conceptual nature of the employed language models support this: semantic similarity reflects a more abstract linguistic prediction that does not necessarily commit to a specific lexical item, whereas surprisal, which predicts specific word identity, maps more formally to prediction error. Future work, however, using more elaborate models should help to better disentangle contributions from these neural populations. That said, the measures show a clear dissociation in their influence on bottom up sensory processing. Furthermore, the dissociation in the timing of these processes is consistent with intuitions behind predictive coding.

In conclusion, our results support predictive coding accounts of how top-down prior knowledge influences bottom up sensory processing during naturalistic speech comprehension.

